# An Antibody Cocktail-Based Immunoaffinity-LC-MS Method Enabled Ultra-Sensitive and Robust Quantification of Circulating Proinsulin Proteoforms and C-peptide

**DOI:** 10.1101/2025.06.02.657397

**Authors:** Qingqing Shen, Wang Cao, Ming Zhang, Tai-Tu Lin, Gabriela S. F. Monaco, Lorenz Nierves, Tujin Shi, Cornelia Boeser, Scott Peterman, Carmella Evans-Molina, Emily K. Sims, Wei-Jun Qian, Jun Qu

**Affiliations:** The Department of Pharmaceutical Sciences, University at Buffalo, Buffalo, NY 14214; New York State Center of Excellence in Bioinformatics and Life Sciences, Buffalo, NY 14203; Biological Sciences Division, Pacific Northwest National Laboratory, Richland, WA 99352; Department of Pediatrics, Indiana University School of Medicine, Indianapolis, IN 46202; The Center for Diabetes & Metabolic Diseases, Indiana University School of Medicine, Indianapolis, IN 46202; The Herman B. Wells Center for Pediatric Research, Indiana University School of Medicine, Indianapolis, IN 46202; Roudebush Veteran’s Affairs Medical Center, Indianapolis, IN 46202; Thermo Fisher Scientific, San Jose, CA 95134

**Author notes:** The authors contributed equally to this work (Qingqing Shen, Wang Cao). Corresponding Authors: Wei-Jun Qian, Biological Sciences Division, Pacific Northwest National Laboratory, Richland, Washington 99354, United States., Jun Qu, Department of Pharmaceutical Sciences, University at Buffalo, Buffalo, New York 14214, United States; New York State Center of Excellence in Bioinformatics and Life Sciences, Buffalo, New York 14203, United States.

## Abstract

Accurately measuring circulating proinsulin proteoforms is crucial for clinical investigation of diabetes, but was previously not feasible owing to limited assay specificity/sensitivity. Here we devised a highly sensitive LC-MS-based strategy to quantify intact proinsulin, des-31,32 and des-64,65 proinsulin, and C-peptide in circulation. The method involves: i)quantitative, robust affinity capture using an optimized antibody cocktail, eliminating the severe quantitative bias across multiple proteomes typically introduced when using a single antibody; ii)Lys-C digestion producing unique signature peptides for each proteoform, and iii)trapping-nano-LC coupled with FAIMS/dCV-MS for an ultra-sensitive analysis. The selective trapping/delivery ensured sensitive/selective analysis of the targets while achieving excellent analytical robustness that is critical for clinical assays, and the FAIMS/dCV substantially reduces baseline noise/interferences, further enhancing S/N. The assay achieved exceptional sensitivity, with serum LOQs of 1.7, 2.3, and 3.6 pg/mL respectively for intact-proinsulin, des-31,32 and des-64,65, representing the first assay capable of sensitively quantifying these major circulating proinsulin proteoforms. We applied this assay to 78 subjects, including autoantibody positive(n=20) and new-onset type-1-diabetes(T1D, n=19) with respective age/sex/BMI-matched controls, enabling the first accurate profiling of proinsulin proteoforms in clinical groups. The assay results demonstrated a clear separation of control and new-onset T1D groups that a parallel total-proinsulin ELISA assay fails to capture. Furthermore, distinct expression patterns in relative abundance ratios among proteoforms were observed across clinical groups. This assay may provide valuable insights into the β-cell functions and the onset/progression of diabetes and other associated conditions. Moreover, the strategy is broadly applicable to targeted measurement of other biomarker proteoforms.

## INTRODUCTION

Insulin, a vital hormone secreted by pancreatic beta cells, plays a central role in the regulation of blood glucose levels and overall metabolic homeostasis. Insulin is initially synthesized as the precursor proinsulin, which is then subjected to sequential processing as illustrated in **Figure 1A**, resulting in the formation of two major intermediate proteoforms: des-31,32 proinsulin and des-64,65 proinsulin, which eventually lead to the production of mature insulin and C-peptide. Recent studies have found that defective proinsulin processing is observed in both type 1 diabetes (T1D) and type 2 diabetes (T2D), as well as obesity.^1–5^ Interestingly, the levels of proinsulin and the relative levels between different proinsulin proteoforms in circulation (e.g., intact proinsulin, des-31,32, des-64,65) and C-peptide provide potential for stratifying diabetes risk, distinguishing diabetes types and endotypes, monitoring the progression of the disease, and evaluating the efficacy of clinical interventions.^6–8^ To fully empower the clinical and mechanistic utility of these insulin proteoforms in circulation, it is critical to develop reliable clinical assays specific to each of the proteoforms.

**Figure 1.**
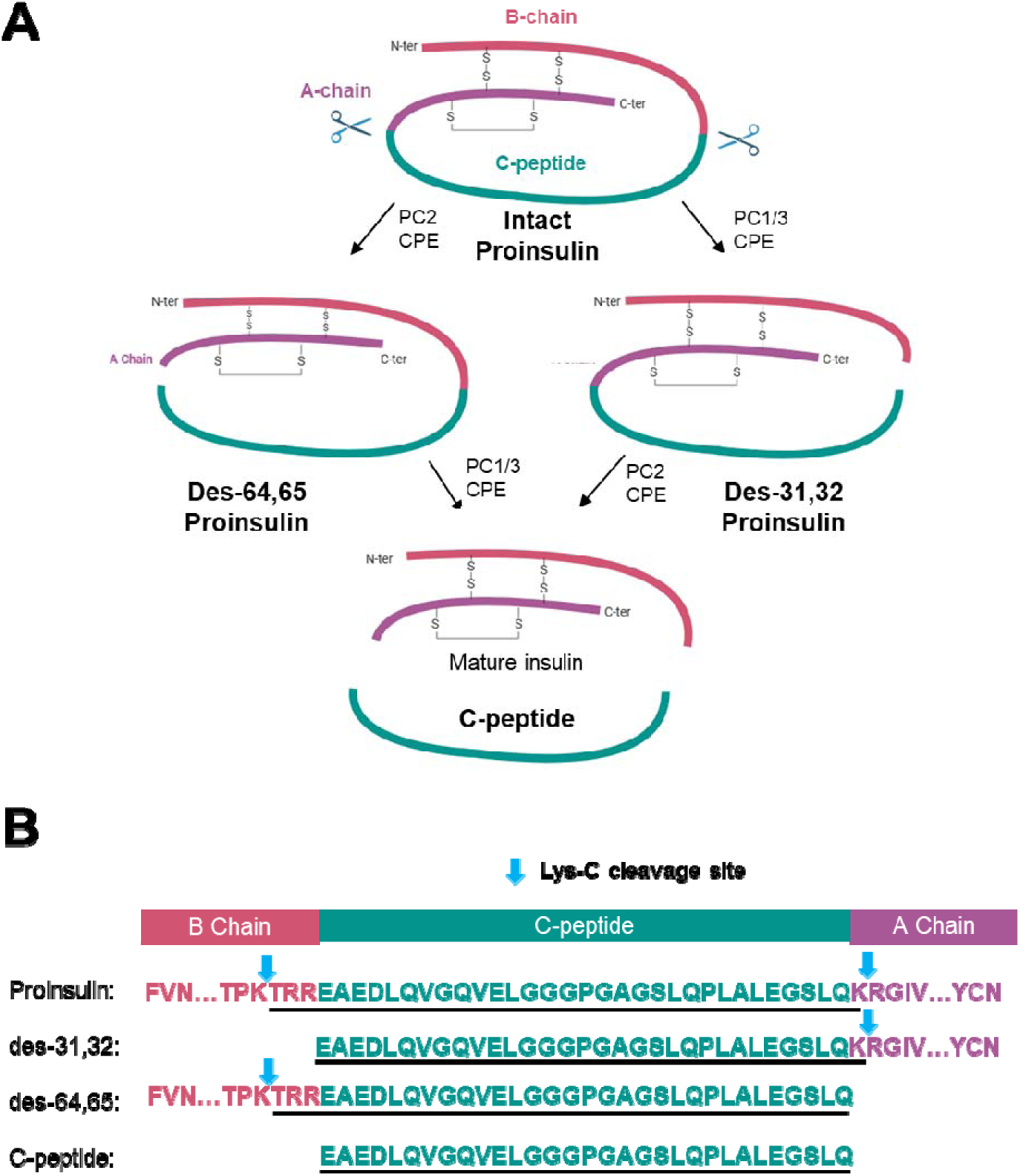
Major proinsulin proteoforms and their specific signature peptides generated by Lys-C digestion. (**A**) Proinsulin processing pathway. Intact proinsulin undergoes selective cleavage at the B-C or A-C junctions by various proteases, forming des-31,32 or des-64,65 proinsulin. These forms are subsequently processed into mature insulin and C-peptide. PC1/3: Prohormone convertase 1/3; PC2: Prohormone convertase 2; CPE: Carboxypeptidase E. (**B**) Signature peptides (underlined) produced by Lys-C digestion, representing each proteoform and C-peptide.

Although quantification of circulating insulin and C-peptide has become a routine practice, measuring proinsulin proteoforms in circulation (i.e., intact proinsulin, des-31,32, and des-64,65) remains a challenge despite many years of efforts. Because these proteoforms have a high degree of similarity in sequences and structures, it is very difficult to distinguish them using conventional immunoassays. Moreover, most antibodies used in existing commercial ELISA kits for proinsulin suffer potential cross-reactivity with the two des-forms of proinsulin,^9^ potentially leading to a substantial overestimation of proinsulin levels in clinical samples.

To overcome this limitation of immunoassays, LC-MS is a promising alternative method for measuring different proinsulin proteoforms due to its high specificity in molecular detection and multiplex capability to quantify several analytes simultaneously. However, to date, a quantitative LC-MS method meeting clinical needs for proinsulin proteoforms quantification, i.e., with sufficient sensitivity, selectivity, and quantitative accuracy, has not been developed. One pioneering work developed an LC-MS method for detection of intact proinsulin and des-31,32 in circulation without interference from other proteoforms, though without quantitative capacity.^10^ A following work attempted semi-quantification of des-31,32 proinsulin.^11^ While the two methods detected proinsulin in some individuals with insulin autoimmune syndrome, where proinsulin levels are substantially higher than other populations including controls,^12^ neither method was sensitive enough to detect proinsulin in control subjects. Another work reported a limit of detection of 50 pg/mL for intact proinsulin, which failed to quantify proinsulin in ∼70% of human plasma samples.^13^ Besides, to our knowledge, detection of des-64,65 proinsulin in circulation has never been achieved in any patient population. Despite the lack of quantitative data, it has been speculated that the levels of des-64,65 could be significantly lower than des-31,32 proinsulin.^14, 15^ Therefore, an LC–MS assay capable of sensitively quantifying all three proinsulin proteoforms alongside C-peptide would be a major advance, particularly for investigating diabetes and other conditions linked to β-cell dysfunction.

To address this need, we devised a novel immunoaffinity LC-MS strategy for absolute quantification of the three main proinsulin proteoforms and C-peptide. Specific signature peptides after Lys-C digestion are strategically selected to permit quantification of each proteoform. By integrating a unique, antibody cocktail-based immunoaffinity capture procedure and a highly sensitive nano-LC-Field Asymmetric Waveform Ion Mobility Spectrometry/differential-Compensation-Voltage (FAIMS/dCV)-MS strategy, limits of quantification (LOQs) at 1.7-3.6 pg/mL were achieved for the three proinsulin proteoforms. This approach represents the first reliable method for quantifying proinsulin and its des-forms in clinical samples.

The method was successfully applied to a cohort of individuals either with T1D or at-risk for T1D, as well as their age/sex/BMI-matched controls, respectively. Our data revealed that the LC-MS assay had superior sensitivity and specificity compared to a regulator-approved, commercially available ELISA assay. Owing to its high sensitivity, the LC-MS-based method confidently quantified intact proinsulin and des-31,32 form in all clinical samples. Notably, this is the first time that the low-level des-64,65 form has been reliably quantified in circulation. We anticipate that the ability to quantify multiple circulating proinsulin proteoforms enables future clinical applications in assessing the utility of these biomarkers for β-cell function, disease progression and/or therapeutic interventions.

## EXPERIMENTAL SECTION

Details for chemical reagents, confirmation of identification of endogenous proteoforms by Orbitrap Astral, quantitative method validation, and data analysis are in the **Supplemental Experimental Section**.

### Protein/peptide standards and chemical reagents

The WHO international proinsulin standard (09/296) was obtained from NIBSC. The C-peptide, des-31,32, and des-64,65 standards and the heavy isotope-labeled peptides were from Biosynth (Staad, Switzerland). The accurate purity of the synthesized standards was rigorously calibrated by quantitative amino acid analysis. The Sulfo-NHS-LC-biotin antibody biotinylation kit was purchased from Thermo Scientific (Waltham, MA). The Dynabeads™ MyOne™ Streptavidin T1 beads were from Invitrogen (Waltham, MA). N-Dodecyl beta-D-maltoside (DDM) was from Sigma-Aldrich (Darmstadt, Germany). TCEP and 2-chloroacetamide were purchased from Thermo Scientific (Waltham, MA). The monoclonal antibodies clone 7F8 and 5E4/3 were obtained from Bio-Rad (Hercules, CA), and clone C-PEP-01 was purchased from Sigma-Aldrich (Darmstadt, Germany). Lys-C enzyme was from New England Biolabs (Ipswich, MA). The human proinsulin ELISA kit (cat# 10-1118-01, Mercodia, Uppsala, Swede), the only such product approved by EU/EEA, UK and Canada. Other reagents are shown in SI.

### Serum samples

The 78 de-identified archived human serum samples were provided by Indiana University. The demographic information of the subjects is summarized **Table S3.** Sample collections were approved by the Indiana University Institutional Review Board. Written informed consent or parental consent and child assent were obtained from all participants before any research participation in accordance with the ethical guidelines of the institution. 200 µL of serum was used for analysis for each sample. Prior to further preparation, the human serum samples were centrifuged at 18,000 g for 30 minutes, and the lipid layer was removed. The pooled equine serum, as the surrogate matrix for calibration and validation, was purchased from Equitech-Bio (Kerrville, TX), and pooled human serum samples for method development were from Innovative Research (Novi, MI).

### Antibody-cocktail-based immunoaffinity capture

To avoid quantitative biases across multiple proinsulin proteoforms associated with using a single pan-antibody, we employed an antibody-cocktail strategy for target protein capture. We screened multiple commercially available anti–C-peptide, anti–insulin, and anti–proinsulin mAbs, testing each individually and in various binary or ternary combinations. For each tested combination, we optimized key conditions such as antibody amounts, antibody-to-bead ratios, and incubation parameters. After extensive trials, three mAbs—5E4/3 (anti–proinsulin), 7F8 (anti–insulin), and C-pep-01 (anti–C-peptide)—were selected because of their superior capture efficiency and uniform recovery across different proteoforms. In the final optimized procedure, each antibody was individually biotinylated in PBS with a 50-fold molar excess of NHS–LC–biotin, using the EZ-Link Sulfo–NHS–LC–Biotinylation Kit and following the manufacturer’s protocol. The reaction was performed in a total volume of 100 µL per 100 µg of mAb. The biotinylated C-pep-01, 7F8, and 5E4/3 were then mixed at a 1:1:1 ratio and stored at 4 °C for up to 30 days. We performed the immunocapture procedure on a KingFisher Flex system (ThermoFisher). For each sample, we used 2 µg each of C-pep-01, 7F8, and 5E4/3 and 0.6 mg of magnetic beads. In each well, 200 µL of serum sample was incubated overnight at 4 °C with the antibody cocktail plus 300 µL of 1× PBST (PBS containing 0.01% Tween-20, pH 7.4) and a protease inhibitor (Sigma-Aldrich, Darmstadt, Germany), with continuous shaking at 300 rpm. The detailed plate setup and KingFisher procedure are provided in the **SI Experimental**. Immediately after elution, the eluates were neutralized with 3 µL of 1 M Tris solution and transferred to low-binding tubes for one-step reduction, alkylation, and rapid Lys-C digestion, as also described in the **SI Experimental**.

### Trapping-nano-LC-FAIMS-MS analysis

An UltiMate 3000 LC system (NCS-3500RS CAP, micro/nano binary pumps, and WPS-3000 TBRS autosampler) was coupled to a FAIMS Pro Duo interface on an Altis Plus triple-quadrupole mass spectrometer (ThermoFisher). The mobile phases were prepared as follows: A=5:95:0.1 (water:acetonitrile:formic acid, v/v/v) and B=88:12:0.1 (water:acetonitrile:formic acid, v/v/v). Samples were introduced from the autosampler via a 5-µL loop to a large-ID trap column (150 µm i.d. × 5 cm, 5 µm, 300 Å C18, CoAnn Technologies) at 20 µL/min and 13% B, enabling rapid, selective retention of peptides while flushing out hydrophilic matrix components. The initial nano flow was set to 750 nL/min at 18% B for 1.0 min. The trap was then switched in-line with the nano column (75 µm i.d. × 10 cm, 1.7 µm, 100 Å C18, CoAnn Technologies), and peptides were backflushed from the trap to column at 750 nL/min. To focus targets at the front end of the column, the gradient was ramped from 18% to 30% B over 1 minute, then to 34.8% B over the next 2.5 minutes at 750 nL/min. The flow rate was subsequently reduced to 250 nL/min, and B increased linearly to 49.2% in 10 minutes, followed by a ramp to 97.0% B over 2.5 minutes. At the 12.5 minutes’ mark, the flow was returned to 750 nL/min and held at 97.0% B for 1 minute. Finally, the column was re-equilibrated at 18.0% B for the remainder of the 15-minute cycle. The trap was switched off column at 8 minutes to prevent hydrophobic components from entering the nano column. Then the trap was flushed in parallel at 20 µL/min with 99% B for 3 minutes to remove residual hydrophobic species, then equilibrated at 13% B for 2 minutes. Both the trap and analytical columns were maintained at 60 °C. Flow path coordination was performed by a zero-dead-volume (ZDV) 6-port valve in a heated column compartment.

For MS analysis, the ESI spray voltage was 2.2 kV, and the temperature of ion transfer tube was 250 °C. The isolation windows were 0.7 Th for Q1 and 1.2 Th for Q3. In the FAIMS device, the carrier gas was set to 3.5 L/min, and both the inner and outer electrodes were heated to 100 °C. RF-lens voltage and collision energy were optimized for each signature peptide (SP) via an on-the-fly orthogonal array strategy.^16^ The FAIMS CV settings were determined according to the FAIMS/dCV method.^17^ MRM transitions and FAIMS CV parameters for the targets are listed in **Table S1**.

To eliminate carryover between samples, a programmed syringe-washing procedure was employed after each injection. The autosampler needle was moved to the home position and washed with 5 µL of mobile phase A, followed by two cycles of drawing and dispensing 5 µL of 80% methanol and mobile phase A, respectively. The syringe was then returned to the home position, ready for the next injection.

## RESULTS AND DISCUSSION

### 1. Development and optimization of the trapping-nanoLC-FAIMS/dCV-MS strategy for proinsulin proteoforms

The objective was to develop a multiplex assay for quantification of the three proinsulin proteoforms (**Figure 1A)** along with C-peptide, which also enables the measurement of their relative ratios—an important factor in understanding diabetes and other related conditions. To achieve this, we leveraged the molecular-level selectivity of LC–MS. Specifically, Lys-C proteolysis was employed to produce a unique signature peptide for each proteoform **(Figure 1B**). Note that the term “proinsulin” refers to “intact proinsulin” throughout the text unless otherwise specified. The three signature peptides reflect the distinct cleavage patterns at the A-C and/or B-C chain junctions of each proteoform, enabling their specific detection. The SRM conditions for these signature peptides are in **Table S1.**

To achieve sensitive and accurate quantification, an antibody-cocktail-based immunoaffinity LC-MS method was developed. The optimized antibody cocktail enabled efficient and reproducible capture of all proinsulin proteoforms while avoiding quantitative biases that can arise when using a single pan-antibody (**Figure 2**). The optimization and characterization of the antibody cocktail-based enrichment are discussed in detail below. The captured proteins undergo a one-step reduction/alkylation followed by a rapid 3-hour Lys-C digestion, which affords high peptide yields with excellent reproducibility. Notably, the inclusion of a non-ionic, MS-compatible detergent DDM during resuspension and digestion significantly improved quantitative reproducibility (**Table S2**), likely due to DDM’s ability to reduce surface adsorption of the low-abundance targets.

**Figure 2.**
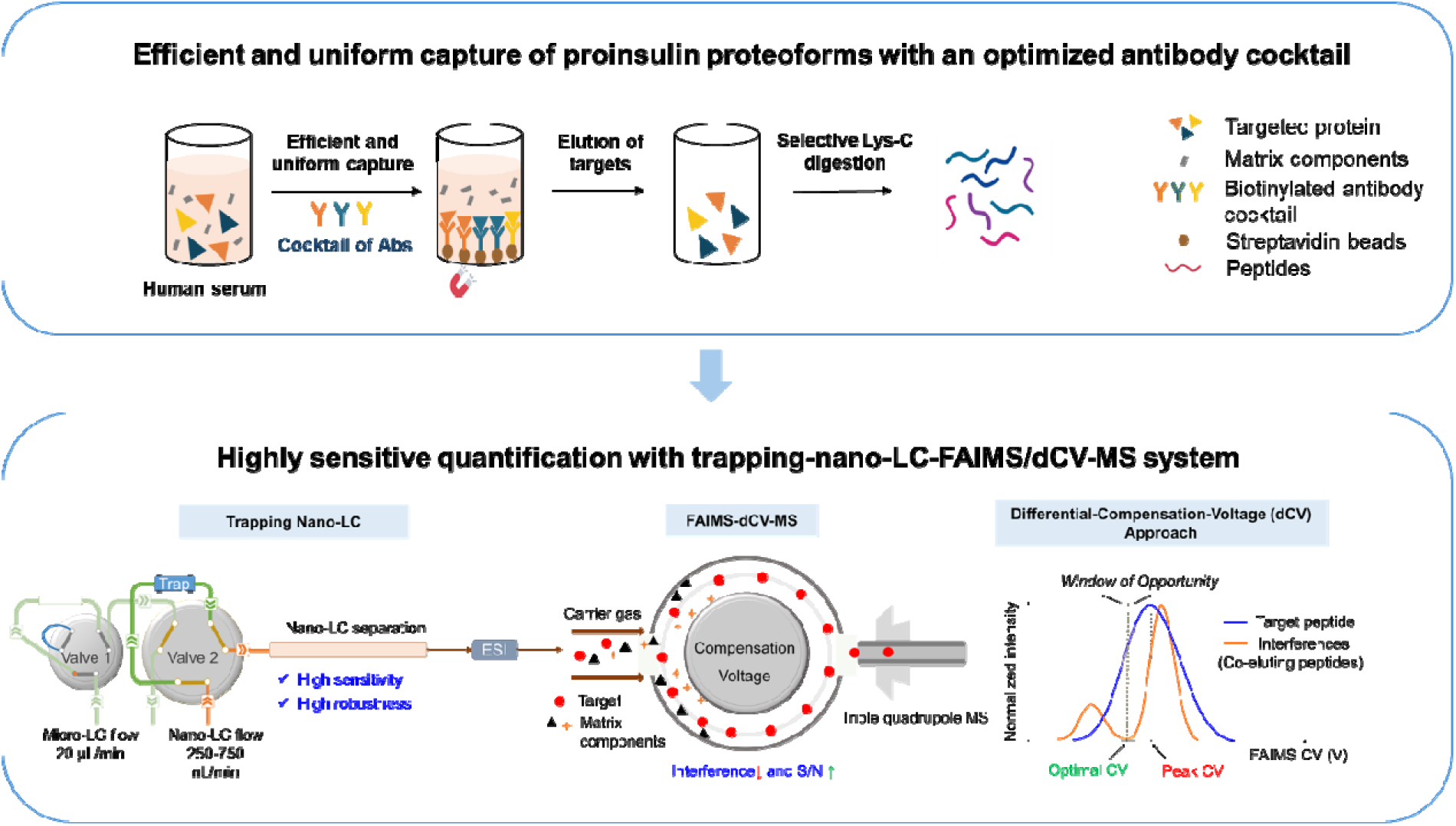
Flowchart illustrating the LC-MS-based strategy for sensitive and robust quantification of proinsulin proteoforms and C-peptide in circulation. Proinsulin proteoforms and C-peptide from serum are efficiently enriched using an optimized antibody cocktail, followed by digestion with Lys-C. The resulting peptides are analyzed using a trapping-nano LC system coupled with FAIMS/dCV-MS, enabling highly sensitive and reproducible measurements.

The resulting digest is then loaded to a trapping-nano-LC-FAIMS/dCV-MS system for sensitive and robust analysis, adapted from the previously described strategy.^17^ The selective trapping/delivery of the targets via a large-ID trap prevented hydrophilic and hydrophobic matrix components from entering the nano-LC-MS system, which improves analytical selectivity, sensitivity, robustness and throughput. Additionally, the large-ID trap permits a homogeneously mixed gradient delivery to the downstream nano-LC column,^18, 19^ contributing to the excellent run-to-run reproducibility discussed below. Additionally, to maintain reasonable throughput while preserving separation quality, a series of flow strategies were developed. A one-minute, rapid focusing gradient effectively transfers targets from the trap to the column without band broadening, followed by a two-stage analytical gradient for efficient separation with high sensitivity. With these approaches, the total cycle time, including sample loading, was controlled within 15 minutes.

Because circulating proinsulin proteoforms are present at extremely low levels, minimizing background noise and matrix interferences—even post-immunoaffinity enrichment—is crucial to achieve desirable sensitivity and selectivity. Here we took advantage of the recently developed FAIMS/dCV approach to substantially reduce the noise levels and therefore further boosting the S/N.^17^ FAIMS’s separation mechanism is orthogonal to both LC and MS, therefore could potentially enhance analytical selectivity. However, traditional FAIMS methods employ the compensation voltage (CV) yielding the highest intensity of each peptide without considering the co-isolated noises, which often results in only marginal improvement of S/N.^17^ Conversely, the FAIMS/dCV identifies the optimal CVs delivering the highest S/N by extensively comparing the CV profiles of the target vs. interferences. **Figure 3A** illustrates the dCV optimization for proinsulin proteoforms in human serum, and the S/N-vs.-CV profiles are shown in **Figure S1**. By minimizing baseline noise and interferences in serum samples (**Figure 3B**), the FAIMS/dCV approach improved S/N by 2–11 times for three proinsulin proteoforms relative to analyses without FAIMS. Moreover, prominent inferences that were not readily resolved chromatographically were eliminated by FAIMS/dCV (asterisks in **Figure 3B**). The optimized FAIMS/dCV parameters are listed in **Table S1.**

**Figure 3.**
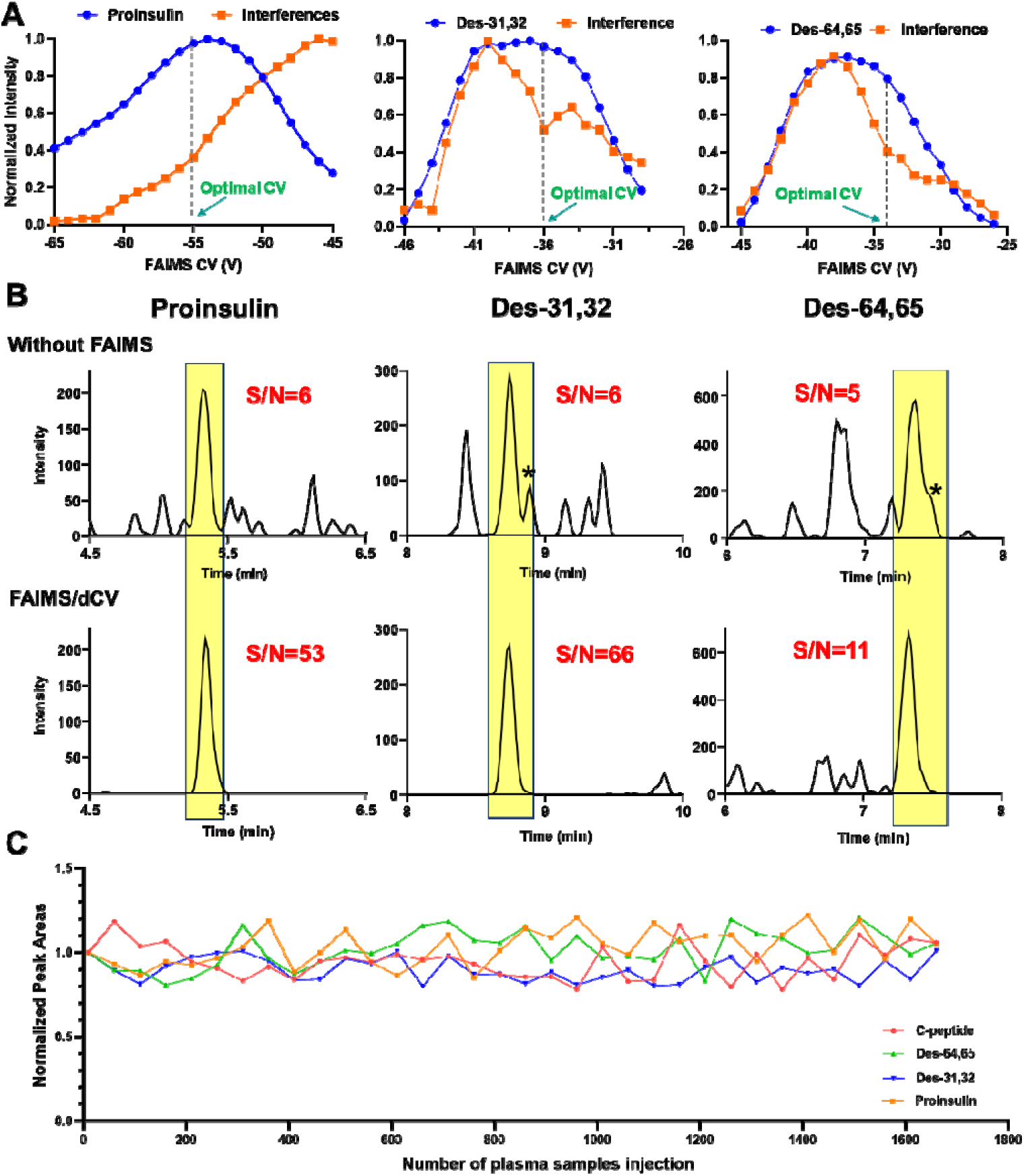
FAIMS/dCV-MS significantly reduces baseline noise and interference, enabling highly sensitive and robust quantification of proinsulin proteoforms. (A) Identification of optimal compensation voltage (CV) by comparing intensity-CV profiles for target proteoforms vs. background noise/interference. **(B)** Representative chromatograms demonstrating the substantial improvement in sensitivity and selectivity achieved with the FAIMS/dCV-MS method for proinsulin, des-31,32, and des-64,65 in pooled human plasma samples. S/N indicates the ratio of signal to baselin noise/interference. The asterisk (*) indicates a prominent interference coeluting with the target peak that is removed by FAIMS/dCV. **(C)** Exceptional robustness of the trapping-nano-LC coupled to FAIMS/dCV-MS, demonstrated by consistent signal intensities in a pooled QC sample injected every 40-100 runs among other serum samples, with no significant signal decrease observed after >1600 injections. These results underscore the suitability of this method for clinical applications.

The analytical strategy yielded exceptional sensitivity, with limits of detections (LODs) of 0.9, 1.2, 1.8 and 2.2 pg/mL respectively for proinsulin, des-31,32, des-64,65, and C-peptide. Assay selectivity was confirmed by a series of extended gradient separations in human serum, with no detectable interferences in any target channel. Additionally, a highly sensitive Orbitrap Astral operating under product ion scan mode further confirmed detection specificity of endogenous proinsulin proteoforms in serum by verifying precursor *m/z*, isotopic distributions, and product ion profiles (**Figure S2**).

This method also displayed excellent reproducibility and robustness. Over two consecutive days, the CV% for signal intensities and retention times for all four targets remained below 7% and 5%, respectively. No carryover was observed, likely due to a rigorous autosampler cleaning protocol (see **Supplemental Experimental**). Furthermore, signal intensities remained stable over 1600 serum injections across two months without needing instrument cleaning (**Figure 3C**). These observations highlight the robustness of the method.

### 2. An antibody cocktail-based immunoenrichment strategy minimizes capturing bias and enables accurate measurements of all proteoforms

A single “pan-antibody” targeting the C-peptide domain, theoretically, should enrich all proinsulin proteoforms, as C-peptide is a common region to all. Nonetheless, our study indicated that using a pan-antibody alone could lead to severe quantitative biases among different proteoforms. As shown in **Figure 4A**, when a popular mAb (C-PEP-01) targeting the C-peptide epitope was employed,^20^ the recoveries of the three proteoforms from a pooled plasma sample differed markedly, 33% for proinsulin, 35% for des-31,32, 63% for des-64,65, and 76% for C-peptide.

**Figure 4.**
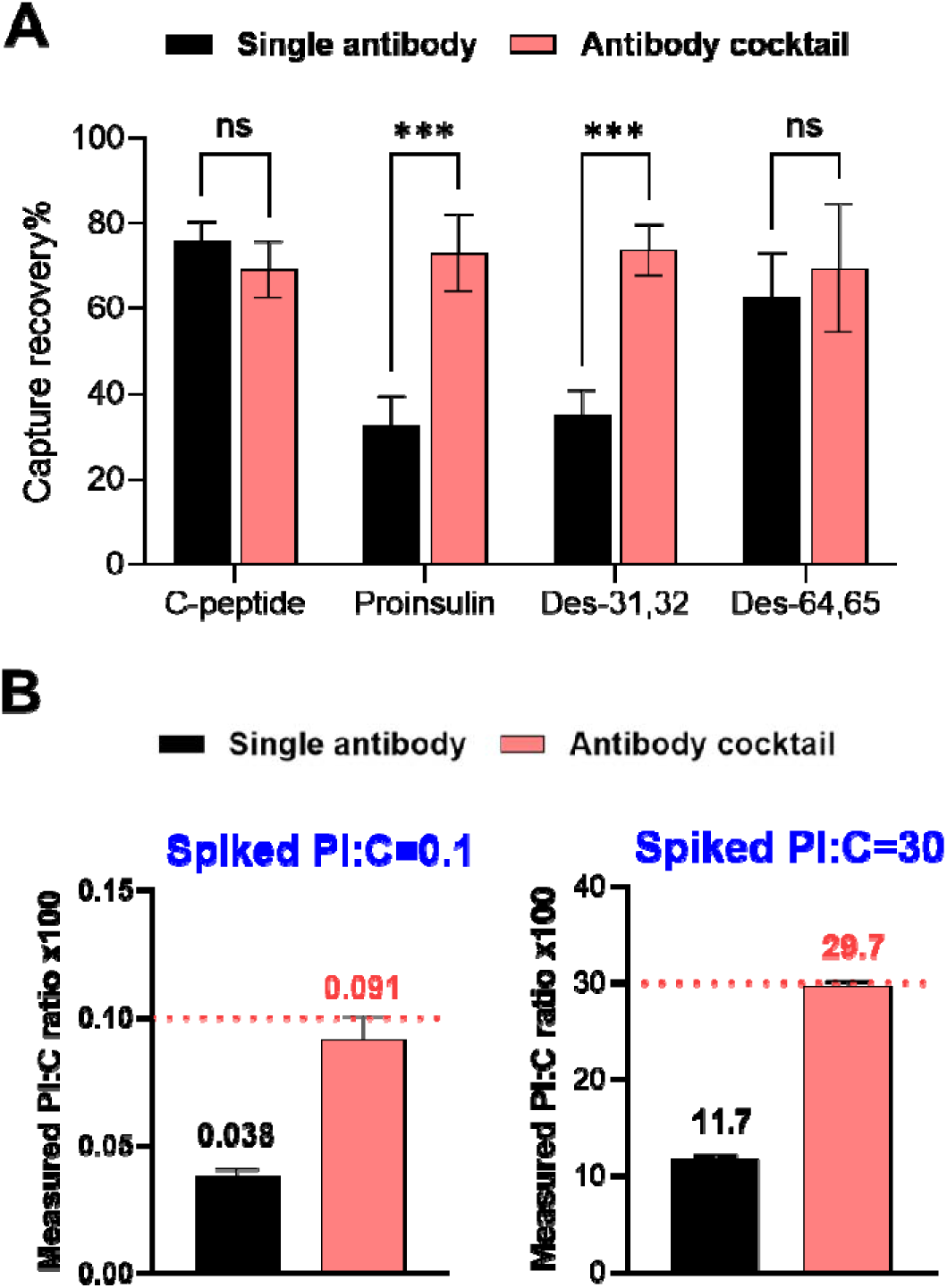
Antibody-cocktail significantly improves accuracy in quantifying proteoform ratios. **(A)** Unequal recovery was observed among different proteoforms when using anti-C-peptide antibody alone, while consistent and high recovery was achieved for each form when using the optimized antibody cocktail. Capture recovery was evaluated in three separate batches, with each value representing the average of two measurements within one batch. **(B)** At both spiked proinsulin to C-peptide ratios in pooled equine serum, there was a severe negative bias in the measured proinsulin to C-peptide ratio when using a single pan anti-C-peptide antibody, while the ratio was nearly maintained when using an antibody cocktail. PI:C ratio: Proinsulin to C-peptide molar ratio×100. Single antibody: 6 μg anti-C-peptid antibody clone C-PEP-01; Antibody cocktail: a mixture of clone C-PEP-01, 5E4/3, and 7F8, with 2 μg each.

Despite extensive optimization of the capture protocol, such as adjusting antibody-to-bead ratios and incubation times, the capturing bias persisted. Moreover, capture efficiencies of individual proteoforms varied considerably across different samples (data not shown). Several other commercially available anti-C-peptide antibodies were also evaluated, all exhibiting similar limitations. Testing multiple anti-proinsulin and anti-insulin antibodies also failed to yield consistent capture of all proteoforms when used alone (data not shown). The discrepancy in recoveries across different proteoforms and different matrices, likely stems from the distinct 3D conformations of individual proteoforms, which affect how the single antibody accesses and binds to targeted epitopes. As a result, different proteoforms exhibit varied capture efficiencies, undermining the accuracy of subsequent measurements.

Such biases could profoundly skew the ratios among the proinsulin proteoforms and C-peptide measured in patient samples, ratios that are critical to assessing for β-cell function.^6, 8^ To demonstrate this risk, we spiked serum with varying molar ratios of proinsulin and C-peptide (0.1 and 30, expressed as a factor of 100), covering the estimated ranges in both controls and T1D subjects.^8, 21^ The spiked samples were immunocaptured with the C-PEP-01 mAb and analyzed by LC-MS. The measured spiked proinsulin-to-C-peptide (PI:C) ratios, respectively 0.038 and 11.7(**Figure 4B**), were >60% lower than the true values.

We hypothesized that such quantitative biases could be alleviated by employing a cocktail of multiple mAbs, each targeting various epitopes on the proteoforms. This approach could provide broader capture coverage and improve the capture efficiency for the proteoforms that are poorly recognized by a single pan-antibody, particularly proinsulin and des-31,32. To test this, we screened various combinations of anti-C-peptide, anti-proinsulin, and anti-insulin antibodies. After extensive trials, an optimal cocktail of three mAbs was identified: 5E4/3 (anti-proinsulin), 7F8 (anti-insulin), and C-PEP-01. Key parameters, such as the amount of each antibody and the antibody-to-bead ratio, were then optimized. The optimized conditions are in the Supplemental Experimental Section.

As shown in **Figure 4A**, this optimized cocktail yielded efficient capture with uniform recoveries across all proteoforms, effectively mitigating the bias observed with the single-antibody strategies. Consequently, when we repeated the spike experiments using the antibody cocktail, the measured proinsulin-to-C-peptide ratios were within 10% of the expected values (**Figure 4B**). These findings confirm that an antibody-cocktail approach can significantly improve both the accuracy and the reproducibility of quantifying multiple proinsulin proteoforms.

### 3. Validation of the quantitative method

As we aim to quantify endogenous targets in human serum, a surrogate matrix that closely resembles human serum is required for calibration and validation to ensure accurate quantification. To identify an appropriate surrogate matrix, we performed parallelism assessments following a previously published protocol.^22^ Briefly, a pooled human serum sample was serially diluted with one of several candidate matrices (guinea pig, equine, and chicken sera), and dilution linearity was evaluated. Among these, equine serum yielded the most linear response: maintaining good linearity for 50-fold dilution (**Figure S3**), whereas other matrices showed inferior linearity.

Then we used equine serum to construct the calibration curves and prepared QC samples for method validation, as detailed in the Supplemental Experimental Section. As shown in **Figure S4**, each target exhibited excellent linearity (R² > 0.99) across the calibration ranges. The validation results are summarized in **Table 1**. The LOQ for C-peptide, proinsulin, des-31,32, and des-64,65 was validated at 8.8, 1.7, 2.3, and 3.6 pg/mL respectively with acceptable accuracy and precision. Across low-, mid-, and high-level validations, the quantitative errors remained <12%, and the coefficient of variation (CV) ranged from 1.0% to 14.3%.

**Table 1.** Validation results for quantification of the four targets in serum.

| Analyte | Linearity range<br>(pg/mL) | QC accuracy (%)/CV% (n=3) |  |  |  |
| --- | --- | --- | --- | --- | --- |
|  |  | <sup>a</sup>LOQ | <sup>b</sup>Low-QC | <sup>b</sup>Mid-QC | <sup>b</sup>High-QC |
| C-peptide | 8.8- 4400 | 97.5/16.6 | 96.0/4.3 | 94.1/4.9 | 107.1/6.7 |
| Proinsulin | 1.7 – 337 | 100.2/9.6 | 97.9/4.2 | 88.8/1.0 | 105.1/3.2 |
| Des-31,32 | 2.3 – 468 | 93.3/9.9 | 98.7/13.8 | 106.3/9.6 | 108.4/6.9 |
| Des-64,65 | 3.6 – 365 | 87.0/1.2 | 106.7/11.6 | 105.6/14.3 | 98.5/3.6 |

<sup>a</sup>The validated LOQs for C-peptide, Proinsulin, Des-31,32 and Des-64,65 are respectively 8.8, 1.7, 2.3 and 3.6 pg/mL in serum.
<sup>b</sup>The levels of High, Mid, and Low QC in serum: C-peptide 3520.0, 440.0, and 22.0pg/mL; Proinsulin at 272.0, 34.0 and 4.3pg/mL; Des-64,65 at 292.0,36.5 and 4.6 pg/mL; Des-31,32 at 372.0, 46.5 and 5.8pg/mL.

### 4. Demonstration of the assay utility in a clinical cohort

To demonstrate the utility of the multiplex assays of proinsulin proteoforms, we applied the method to serum samples from a total of 78 individuals, including a group with very recent onset Stage 3 T1D (within 48 of hospitalization) (n=20), an at-risk group (n=19, individuals considered at-risk for T1D due to being relatives of individuals with T1D subjects and having multiple autoantibodies)^23^, and controls for the new-onset and at-risk groups, respectively, matched by age/sex/BMI. The demographic information of the subjects is in **Table S3**.

All serum samples (200 µL each) were processed and analyzed in a single batch, and the assays’ performance—sensitivity, specificity, accuracy, and precision—was closely monitored by analyzing a group of quality assurance samples, including spiked QC samples, matrix– and solvent-blank samples, and a pooled serum sample. These samples were injected after every 20 injections of patient samples to confirm assay performance. Owing to the unprecedented sensitivity of the assay, proinsulin, des-31,32 proinsulin, and C-peptide were quantified in all subjects, while des-64,65 proinsulin was detected in 80% of the cohort. Box-and-whisker plots of the four targets are shown in **Figure 5A**. For proinsulin, the levels in 75% of the samples are <36 pg/mL, consistent with previous reports indicating that with an LOD of 50 pg/mL, proinsulin was not detectable in most clinical plasma samples.^13^

**Figure 5.**
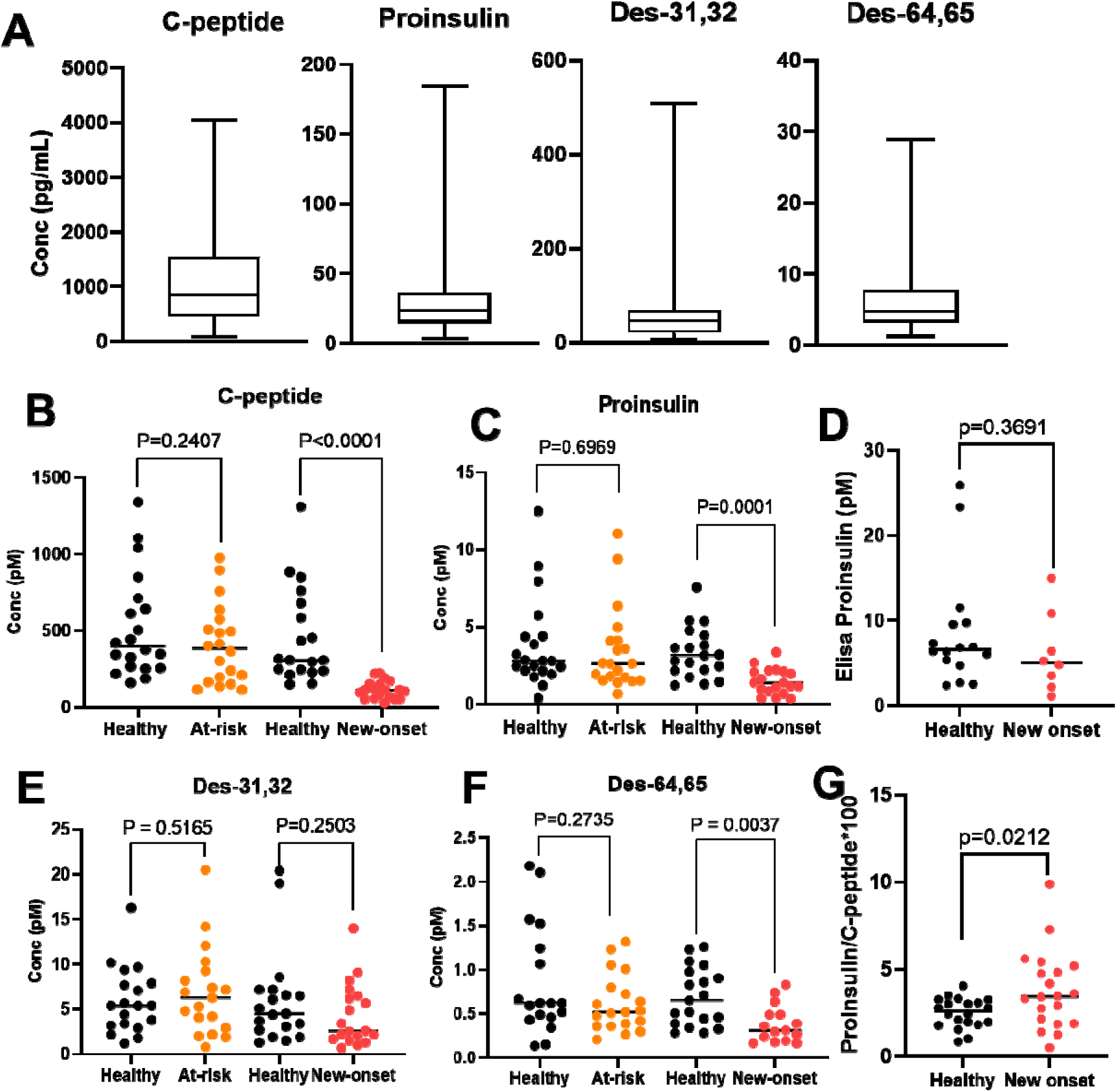
LC-MS-based quantification of proinsulin proteoforms in a clinical cohort of 78 subjects divided into four groups: Controls vs. At-risk for T1D (n=20), Controls vs. new-onset T1D (n=19). **(A)** Box-and-whisker plots summarizing concentrations of the four analytes across all samples. Individual patient levels for **(B)** C-peptide, **(C)** Proinsulin, **(E)** Des-31,32, and **(F)** Des-64,65 is presented as scatter plots with median bars, across the four clinical groups. For comparison, **(D)** shows proinsulin level measured by a parallel total-proinsulin ELISA assay, and **(G)** illustrates the calculated ratios of Proinsulin to C-peptide.

To our knowledge, this is the first study to quantify des-31,32 proinsulin and to detect des-64,65 proinsulin in circulation. While the des-31,32 proinsulin levels in this cohort were comparable to those of proinsulin, des-64,65 proinsulin was present at much lower concentrations, less than one-tenth of the des-31,32 levels. This observation supports previous hypotheses that circulating des-64,65 levels may be significantly lower than des-31,32.^5, 15^

Next, we explored the profiles of various proteoforms among the clinical groups. As shown in **Figure 5B**, the new-onset T1D group displayed significantly decreased C-peptide levels compared to matched controls, consistent with previous findings.^24^ No significant differences in C-peptide levels were observed between at-risk participants and their controls, in line with previous reports. ^25^

Because this assay enables simultaneous quantification of proinsulin, des-31,32 proinsulin, and des-64,65 proinsulin, we could compare their profiles across different clinical groups for the first time. Proinsulin levels in the T1D group were significantly lower than in controls (P=0.0001, **Figure 5C**). By comparison, a parallel ELISA analysis using a regulator-approved kit (discussed below) detected proinsulin in only 26 of 40 samples across the two groups and failed to distinguish between the two groups (P=0.3691, **Figure 5D**). Thus, the LC-MS–based proinsulin assay showed superior discriminatory power, clearly separating control and T1D participants (**Figure 5C**). Des-31,32 proinsulin levels showed no statistically significant difference between the T1D and control groups (P=0.2503, **Figure 5E**). In contrast, des-64,65 proinsulin levels were significantly lower in the new-onset group (P=0.0037, **Figure 5F**), suggesting altered proinsulin processing in T1D and highlighting the importance of quantifying each proteoform. Across all targets, no significant differences were observed between the at-risk group and its controls (**Figure 5**).

Because this assay permits unbiased, simultaneous quantification of proinsulin proteoforms and C-peptide, we can also examine the ratios among these targets, which may have profound clinical implications in diabetes.^7, 26^ For example, the intact proinsulin to C-peptide ratios were significantly different between the new-onset T1D and control groups (P= 0.0212, **Figure 5G**), which agrees with a previous publication.^8^

### 5. Comparison of LC-MS and Proinsulin ELISA Results

To compare the performance of LC-MS and ELISA, we also analyzed the clinical cohort using the only regulator-approved Mercodia proinsulin ELISA kit, which has received approval in the EU/EEA, UK, and Canada and widely employed in clinical analysis.^27^ The ELISA method was rigorously validated following the manufacturer’s instructions. With a limit of detection (LOD) of 16 pg/mL, the ELISA kit detected proinsulin in 59 out of 79 subjects. In contrast, the LC-MS assay, with a significantly lower limit of quantification (LOQ) of 1.7 pg/mL, successfully quantified proinsulin in all samples.

Since the ELISA assay does not distinguish intact proinsulin from its des-31,32 and des-64,65 proteoforms,^28^ we performed two comparisons using the subset of 59 samples that were quantifiable by ELISA. First, ELISA-measured proinsulin levels were directly compared with LC-MS–measured proinsulin levels, revealing a modest correlation (R²=0.675) and a substantial quantitative discrepancy, with ELISA values substantially exceeding LC-MS values (slope=2.6, **Figure 6A**). Second, ELISA results were compared with the sum of all proinsulin proteoforms quantified by LC-MS. This comparison yielded a markedly improved correlation(R²=0.894) and more consistent quantitative values between the two methods (slope=0.892, **Figure 6B**).

**Figure 6.**
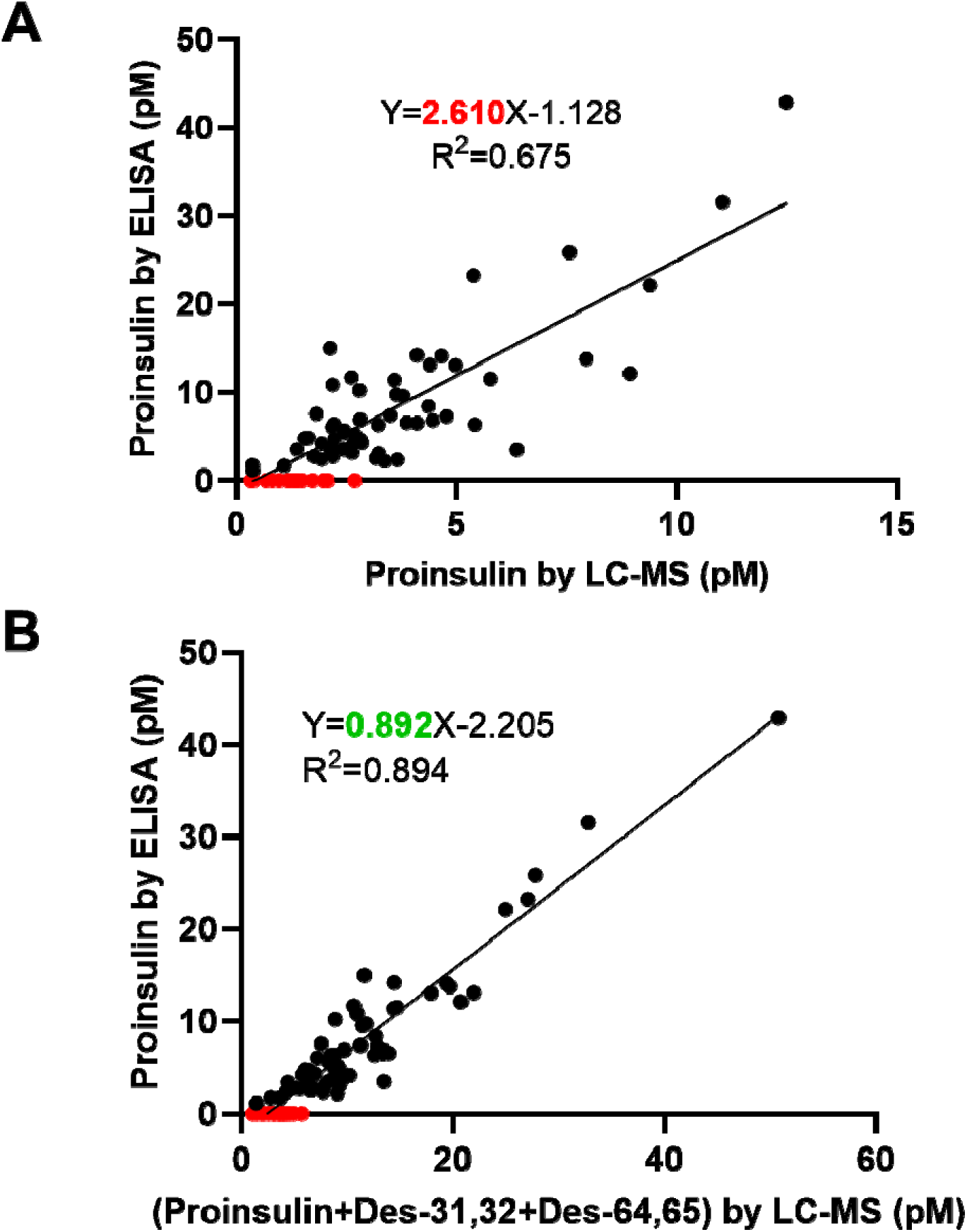
Comparison between LC-MS and ELISA assay for measuring proinsulin in the clinical cohort. **(A)** Correlation between proinsulin levels measured by ELISA and those measured by LC-MS. **(B)** Correlation between total proinsulin levels (the sum of intact proinsulin, des-31,32, and des-64,65) measured by LC–MS and the proinsulin levels measured by ELISA. Red dots represent samples quantifiable by LC-MS but not detectable by the ELISA assay.

These findings confirm previous speculation that the ELISA kit does not differentiate among proinsulin proteoforms, and therefore its reported values likely reflect the total concentration of all proinsulin species rather than proinsulin alone.^28^ Additionally, the results underscore the reliability of LC-MS for accurate and selective quantification of individual proinsulin proteoforms, highlighting its advantage over ELISA in resolving distinct molecular forms.

## CONCLUSIONS

We developed an LC-MS–based assay capable of simultaneously quantifying the three proinsulin proteoforms and C-peptide in human serum with high sensitivity and specificity. Our method employs an antibody cocktail for immunoaffinity enrichment of the four targets, a Lys-C digestion to generate unique signature peptides, and a trapping-nano-LC-FAIMS/dCV-MS to achieve high sensitivity, selectivity and robustness. Compared to the substantial quantitative biases observed with a single “pan-antibody”, the optimized antibody cocktail delivered uniform recovery across all proteoforms and permitted accurate quantification of all proteoforms.^28^

To enable robust LC-MS analyses, we used a relatively large-ID trap for reproducible gradient delivery to the nano-column and rapid sample loading. Moreover, the selective trapping/delivery approach effectively excluded most hydrophobic and hydrophilic matrix compounds, thus maintaining exceptional operational robustness that is on par or beyond that of conventional-flow LC-MS. With an optimized 15-minute cycle, the method holds a practical throughput for clinical applications. Additionally, we leveraged a novel FAIMS/dCV method to significantly reduce baseline noise and matrix interference, thereby further improving overall S/N ratios and pushing the LOQ down to 1.7–3.6pg/mL for all proinsulin proteoforms.

The utility of this assay was demonstrated by analyzing the proinsulin proteoforms in clinical serum samples from 80 subjects with either new-onset T1D or at-risk for T1D, as well as their respective matched controls. For the first time, specific quantification of proinsulin, des-31,32 proinsulin, and des-64,65 proinsulin was achieved at high sensitivity and selectivity in clinical serum samples. The high sensitivity of the LC-MS assay permitted quantification of proinsulin and des-31,32 proinsulin in all clinical samples, while a regulator-approved proinsulin ELISA kit only quantified proinsulin in 74% of samples. Finally, while our findings confirmed that proinsulin-to-C-peptide ratios are higher in T1D subjects compared to controls, the exact significance of different proinsulin proteoforms and their relative ratios in type 1 diabetes is warranted for more extensive follow up studies with clinical samples or longitudinal samples.

We recognize that a “signature peptide” is not intended to target a single proteoform exclusively. Given the complexity of protein processing, variations including truncated forms might produce proteoforms that contain the same signature peptide. Instead, assays based on “signature peptide” will measure a class of proteoforms that share the specific target peptide.^29^ This principle applies to any quantification method that utilizes a unique peptide sequence or epitope, such as LC-MS assays and immunoassays. For proinsulin processing, the intact, des-31,32 and des-64, 65 proinsulin represent three distinct classes of proteoforms, characterized by specific differences at the junctions of B-chain, C-peptide, and A-chain due to enzymatic activity (**Figure 1B**). The signature peptides designed in our assay effectively capture these differences, thus representing each class of proteoforms.

In conclusion, this highly sensitive and selective LC-MS assay provides a robust analytical tool for detailed profiling of multiple proinsulin proteoforms in circulation, and for enabling clinical applications aiming at elucidating the value of these biomarkers in predicting pancreatic beta cell stress, diabetes progression, and responses to therapies. Moreover, our technical approach can be adapted to quantify other critical proteoforms, broadening the potential impact on biomarker discovery and clinical research.

## ACKNOWLEDGEMENTS

This work was supported by NIH grants U01 DK137113 and R01 DK135081, and a Center of Protein Therapeutics consortium grant. Data presented in this work were obtained via the University at Buffalo Center for Proteomics and Bioanalysis.

## Notes

### Competing Interest Statement

The authors have declared no competing interest.

