## Supplemental Information for "An Antibody Cocktail-Based Immunoaffinity-LC-MS Method Enabled Ultra-Sensitive and Robust Quantification of Circulating Proinsulin Proteoforms and C-peptide"

* Corresponding Authors:

Wei-Jun Qian

**Table** **of** **Contents**

### **Supplemental Experimental Section**

**Chemical reagents.** Phosphate-buffered saline (PBS, 10×) was obtained from Corning (Glendale, AZ). Tris was purchased from MP Biomedicals (Irvine, CA). Protease inhibition was achieved using cOmplete™ Protease Inhibitor Cocktail, EDTA-Free, Mini tablets (S	igma-Aldrich, Darmstadt, Germany). Endoproteinase LysC (20 µg) was acquired from Creative Biolabs (Shirley, NY). The detergent n-dodecyl β-D-maltoside (≥98% by GC analysis) was purchased from Sigma-Aldrich (Darmstadt, Germany). Tris(2-carboxyethyl)phosphine hydrochloride (TCEP·HCl) and 2-chloroacetamide (CAA, 98%) were obtained from Thermo Scientific (Waltham, MA).Hydrochloric acid (HCl) was obtained from Fisher Chemical(Pittsburgh, PA). Sample desalting was performed using Zeba Spin Desalting Columns (7k MWCO, 2 mL; Thermo Scientific), and 1.5-mL Protein LoBind Tubes were purchased from Eppendorf (Enfield, CT).

**Kingfisher procedure for enrichment of the targets with an optimized antibody cocktail.** The immunoenrichment process utilized eight 96-well plates arranged as follows:

- Plate 1: 96-well magnetic tip comb.
- Plate 2: Each well contains 60 µL of magnetic beads mixed with 140 µL of 1× PBST (phosphate-buffered saline, pH 7.4, containing 0.01% Tween-20).
- Plate 3: 1 mL of 1× PBST for the initial bead wash.
- Plate 4: Each well holds 200 µL of centrifuged serum sample, 6 µg of the antibody cocktail, and 300 µL of 1× PBST.
- Plates 5 & 6: Each well contains 400 µL of 1× PBST for washing the bound mAb–target complexes.
- Plate 7: 400 µL of 1× PBS for a final wash.
- Plate 8: 100 µL of 25 mM HCl for eluting the captured target proteins.

The KingFisher operation begins by retrieving the plastic comb from Plate 1 and immersing it in Plate 2, where the beads are mixed for 5 minutes. The beads are then washed in Plate 3, transferred to Plate 4 for a 1-hour antigen–antibody binding step, and subsequently washed in Plates 5, 6, and 7 for 5 minutes each. Finally, the target proteins are eluted in Plate 8 with 100 µL of 25 mM HCl. Immediately after elution, the eluates are neutralized with 3 µL of 1 M Tris solution and transferred to low-binding tubes for protein digestion and LC–MS.

**One-step reduction and alkylation, and Lys-C digestion.** The neutralized eluates are evaporated in a SpeedVac and reconstituted in 13 µL of 0.002% DDM, followed by 5 minutes each of ultrasonication and vortex mixing. Next, 1 µL of 100 mM TCEP and 2 µL of 200 mM 2-chloroacetamide are added, and the sample is heated at 95 °C for 10 minutes. After cooling to room temperature, 1 µL of 0.1 µg/µL Lys-C is added, and the mixture is incubated at 37 °C for 3 hours with gentle shaking (280 rpm). The digestion is quenched by adding 1 µL of 13% formic acid and 2 µL of 10 ng/mL internal standard mix, then centrifuging at 18,000 g for 30 minutes. Finally, a 5-µL aliquot of the supernatant is injected into the trapping–nanoLC–FAIMS–MS system.

**Characterization of proinsulin proteoforms in serum samples by high-resolution Orbitrap Astral mass spectrometry.**

To confirm the identity of proinsulin and des-forms in serum samples, we performed parallel MS1 full-scan and product-ion scans on an Orbitrap Astral mass spectrometer. The Orbitrap analyzer acquired MS1 scans at 240,000 resolution (full width at half maximum) across an 800–1200 m/z range, with a 50 ms maximum injection time and an RF lens setting of 50%. Using the monoisotopic precursor masses specified in the mass table (proinsulin at m/z 890.972 and des-31,32 proinsulin at m/z 1049.877), we collected MS2 spectra in the Astral analyzer over 250–1700 m/z, with a 100% AGC target, a maximum injection time of 10 ms, a 30% HCD collision energy, and an isolation window of 1.6 Th.

**Data analysis.** Raw data were processed using the Skyline software (University of Washington, UW, USA), and statistical analysis was performed using GraphPad Prism 10.

**Quantitative method development and validation**.

To construct calibration curves, standards of C-peptide, Proinsulin, Des-31,32, and Des-64,65, with actual purity calibrated by quantitative amino acid analysis, were spiked into pooled equine serum and then serially diluted with pooled equine serum to concentrations ranging from 8.8 to 4400pg/mL, 1.7 to 340 pg/mL, and 2.3 to 460 pg/mL for des-31,32 proinsulin and 3.6 to 360 pg/mL for des-64,65 proinsulin. Six-point calibration curves were prepared for each analyte. All calibration samples were enriched and digested following the experimental protocols. After digestion, the mixture of four SIL-peptide IS was added. Quality control (QC) samples at three concentration levels (High, Mid, and Low) were prepared by spiking standard proteins into equine serum: C-peptide at 3520.0, 440.0, and 22.0 pg/mL; proinsulin at 272.0, 34.0, and 4.3 pg/mL; des-64,65 proinsulin at 292.0, 36.5, and 4.6 pg/mL; and des-31,32 proinsulin at 372.0, 46.5, and 5.8 pg/mL. The precision of the assay was calculated by triplicate preparation of the spiked QC samples, and the coefficient of variation (CV%) of the replicate measurements was calculated to determine the variability. The quantitative accuracies of the QC samples at each concentration level were calculated against the nominal concentrations. For each batch, the calibration regression coefficient of determination (i.e., r^2^) was required to be >0.98. The measured signals for the LLOQ calibrator should be within 20% of the regression line, and all other calibrators should be within 15% of the regression line. The observed concentrations of three levels of QC samples need to be within 20% of their respective nominal concentrations.

### **Table S1. MRM transitions and FAIMS/dCV parameters of the signature peptides**

| Peptide | *Precursor  m/z | *Product  m/z | CE  (V) | RF  (V) | FAIMS CV (V) |
| --- | --- | --- | --- | --- | --- |
| TRREAEDLQVGQVELGGGPGAGSLQPLALEGSLQK (+4) | 891.490 | 528.308 | 24 | 110 | -55 |
|  | 891.490 | 1055.61 | 24 | 110 | -55 |
|  | **891.490** | **1253.636** | **24** | **110** | **-55** |
| TRREAEDLQVGQVELGGGPGAGSLQPLALEGSLQK (heavy) (+4) | 893.475 | 532.316 | 24 | 110 | -55 |
|  | 893.475 | 1063.624 | 24 | 110 | -55 |
|  | **893.475** | **1253.636** | **24** | **110** | **-55** |
| EAEDLQVGQVELGGGPGAGSLQPLALEGSLQ (+3) | 1007.767 | 854.947 | 24 | 124 | -40 |
|  | **1007.767** | **927.515** | **24** | **124** | **-40** |
|  | 1007.767 | 1312.711 | 24 | 124 | -40 |
| EAEDLQVGQVELGGGPGAGSLQPLALEGSLQ (heavy) (+3) | 1010.083 | 858.455 | 24 | 124 | -40 |
|  | **1010.083** | **934.532** | **24** | **124** | **-40** |
|  | 1010.083 | 1319.728 | 24 | 124 | -40 |
| EAEDLQVGQVELGGGPGAGSLQPLALEGSLQK (+3) | **1050.491** | **1055.61** | **34** | **249** | **-36** |
|  | 1050.491 | 1089.592 | 34 | 249 | -36 |
|  | 1050.491 | 1182.132 | 34 | 249 | -36 |
| EAEDLQVGQVELGGGPGAGSLQPLALEGSLQK (heavy) (+3) | **1053.139** | **1063.624** | **34** | **249** | **-36** |
|  | 1053.139 | 1093.599 | 34 | 249 | -36 |
|  | 1053.139 | 1186.139 | 34 | 249 | -36 |
| TRREAEDLQVGQVELGGGPGAGSLQPLALEGSLQ (+3) | 1145.593 | 599.812 | 41 | 160 | -34 |
|  | **1145.593** | **1253.636** | **41** | **160** | **-34** |
|  | 1145.593 | 1394.223 | 41 | 160 | -34 |
| TRREAEDLQVGQVELGGGPGAGSLQPLALEGSLQ (heavy) (+3) | 1147.909 | 599.812 | 41 | 160 | -34 |
|  | **1147.909** | **1253.636** | **41** | **160** | **-34** |
|  | 1147.909 | 1394.223 | 41 | 160 | -34 |

*Transitions in bold numbers are for quantification. The other two transitions are for qualification and confirmation.

### **Table S2. Inclusion of DDM for resuspension and digestion significantly reduced quantification variation**

| **Group** | **CV% for three preparation replicates (n=3)** | | |
| --- | --- | --- | --- |
|  | **C-pep** | **Proinsulin** | **Des-31,32** |
| No DDM | 10.8% | 17.8% | 13.7% |
| DDM at 0.002% | 1.5% | 6.0% | 2.4% |

### **Table S3. The demographic characteristics of the four groups of clinical samples**

|  | At-risk | Controls | New onset T1D | Controls |
| --- | --- | --- | --- | --- |
| Gender: Female/Male, N | 6/14 | 6/14 | 7/12 | 7/12 |
| Age (yr): mean (SD) | 13.4 (10.0) | 13.5 (9.8) | 12.3 (5.9) | 12.4 (6.0) |
| BMI (kg/m^2^): mean (SD) | 20.2 (6.6) | 20.0 (5.7) | 19.4 (4.5) | 20.0 (4.6) |

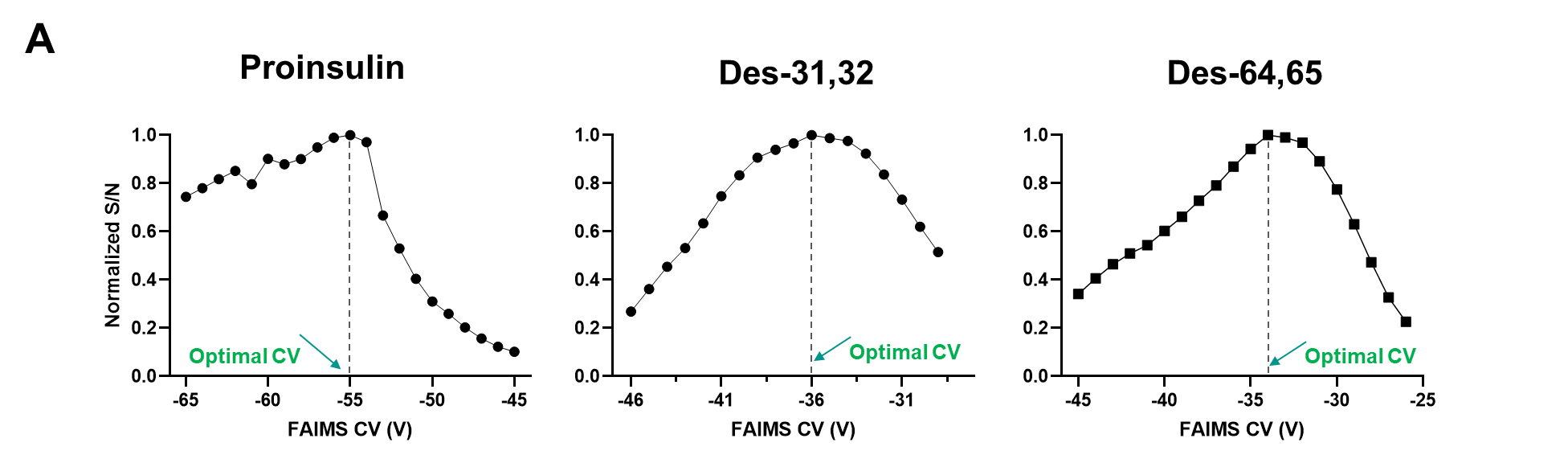

### **Figure S1：The S/N vs. CV profiles for the optimization of the three proinsulin proteoforms in a pooled human serum.**

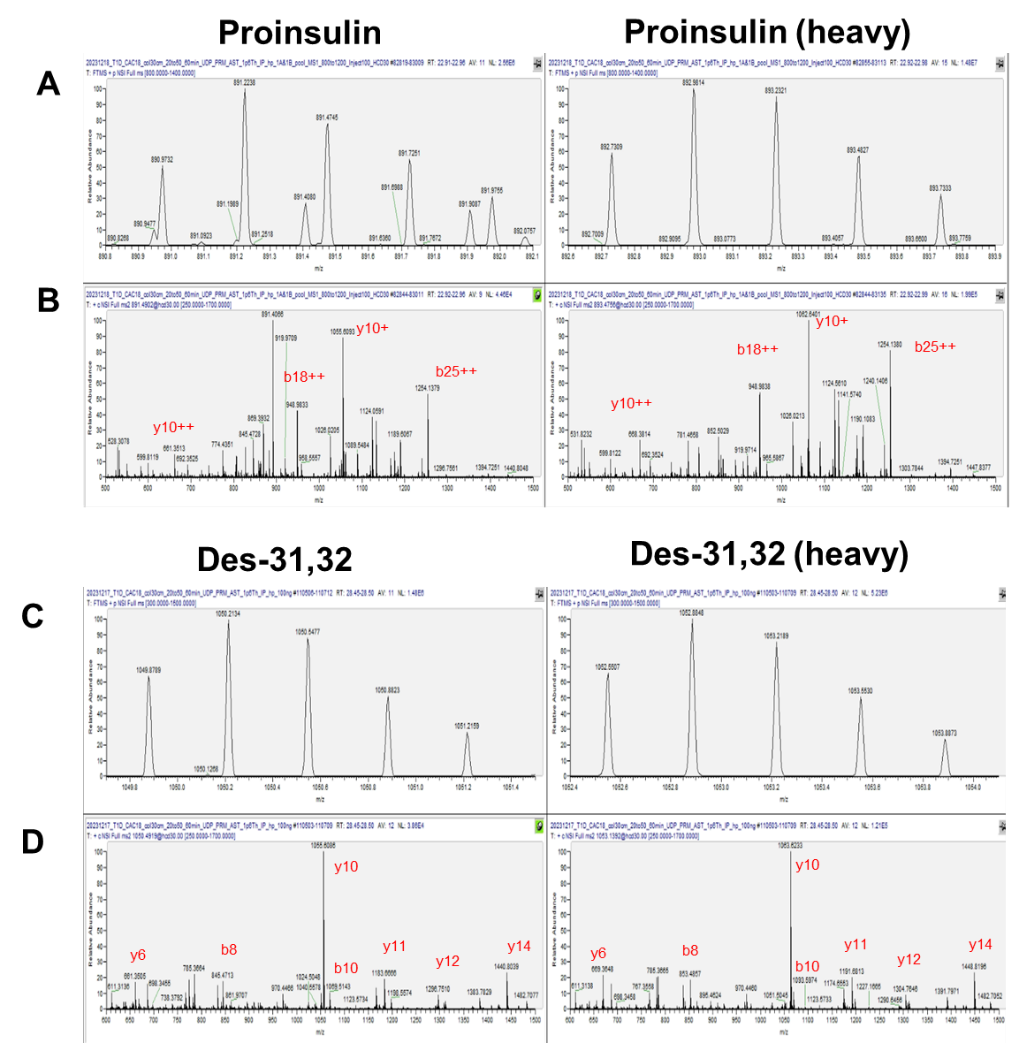

### **Figure S2.** **Verification of specificity for detecting proinsulin and des-31,32 in human serum using a high-sensitivity Orbitrap Astral MS. Endogenous signals were compared with a synthetic heavy-labeled standard.** (**A**) The isotope distribution envelope of proinsulin in serum vs. the spiked standard. (**B**) Astral-generated product ion spectra of proinsulin in serum vs. the spiked standard. (**C**) The isotope distribution envelope of des-31,32 in serum vs. the spiked standard. (**D**) Astral-generated product ion spectra of des-31,32 in serum vs. the spiked standard.

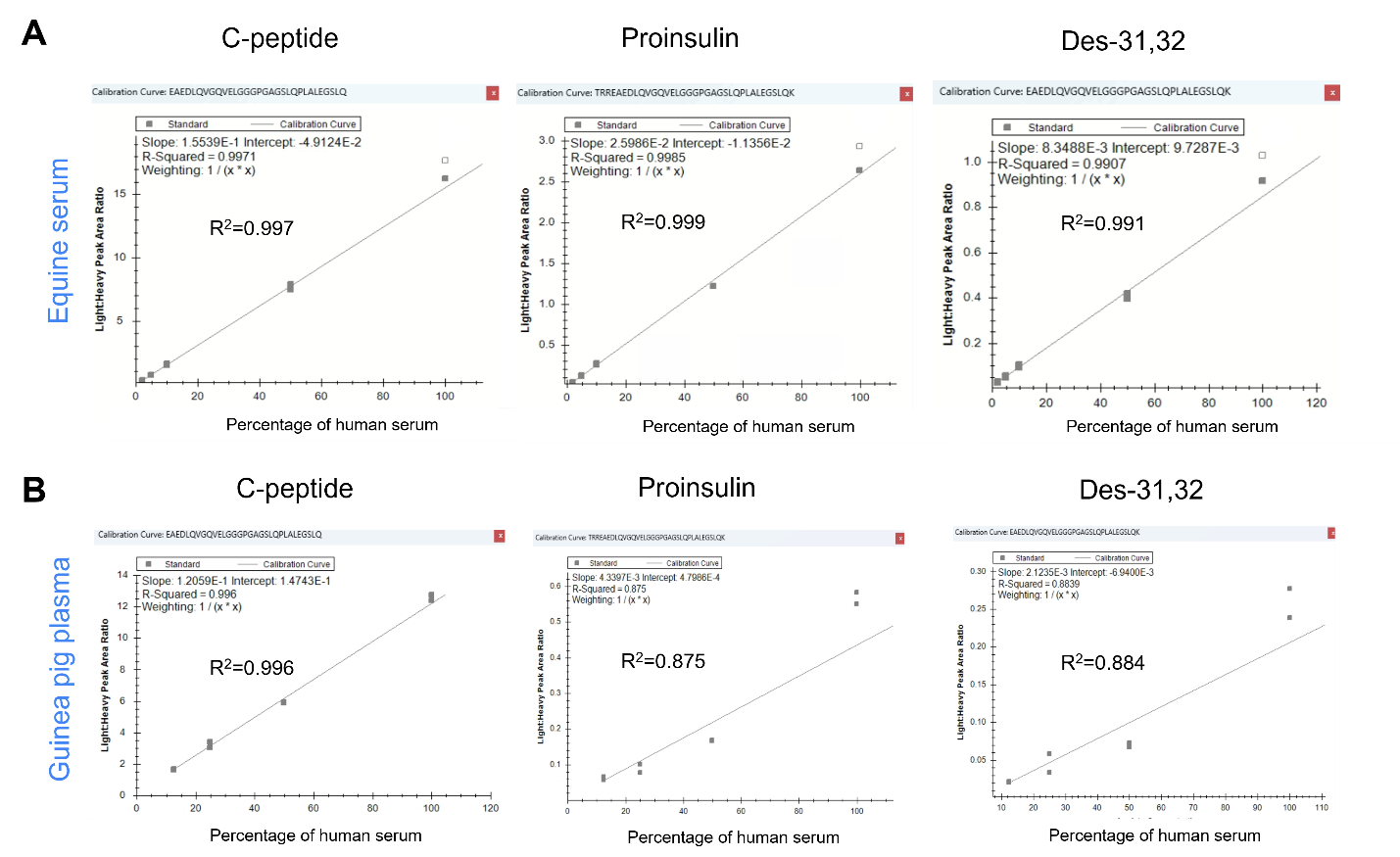

### **Figure S3. Parallelism evaluation via dilution linearity tests. (A)** A pooled human serum sample was serially diluted with equine serum, and the targets were measured experimentally. **(B)** As a comparison, a pooled human serum sample was serially diluted with guinea pig serum, exemplifying that other surrogate matrices we evaluated (in this case, guinea pig serum) resulted in inferior linearity than equine serum.

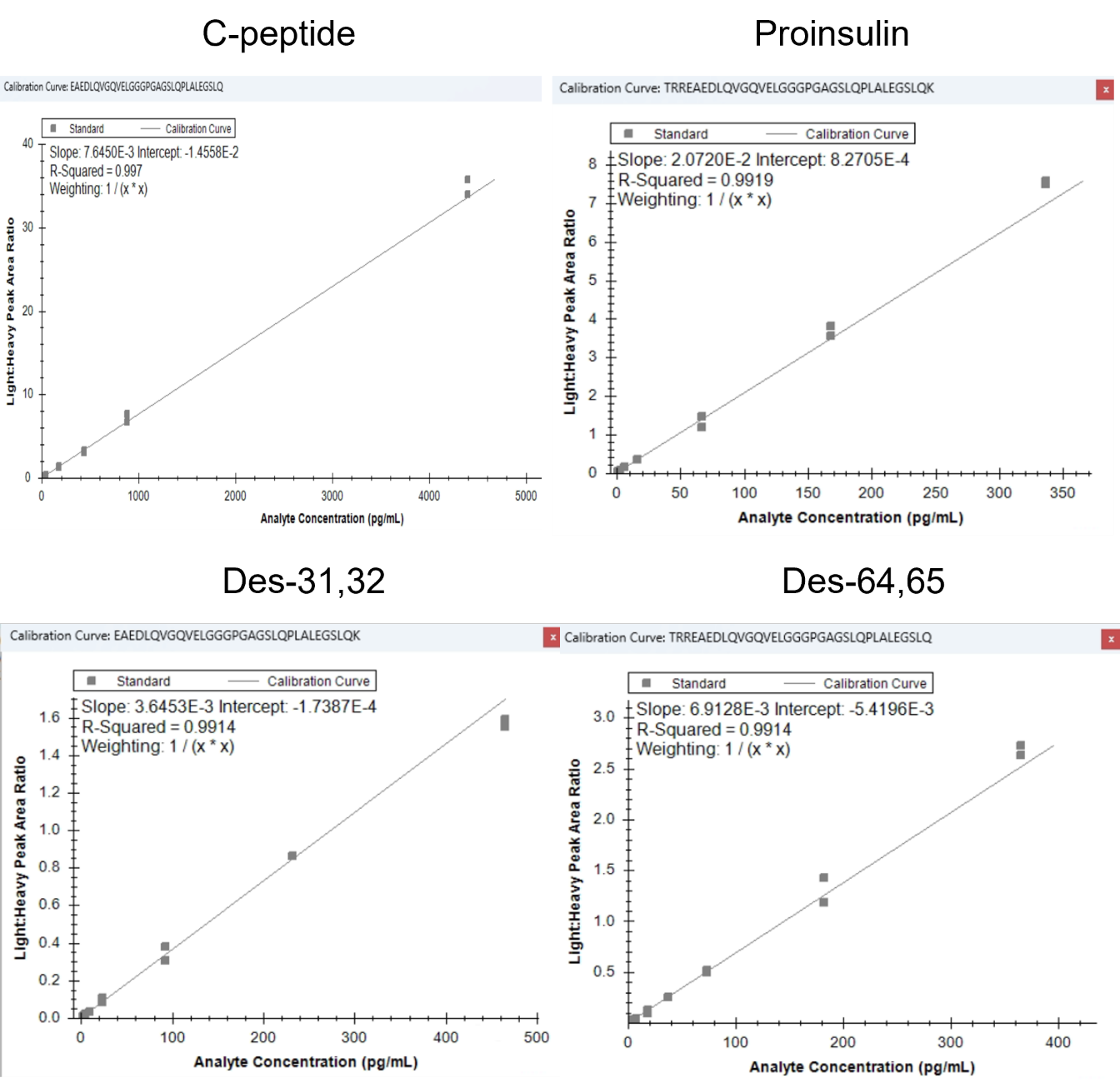

### **Figure S4. Representative calibration curves for the four targets.** The range of concentrations are: C-peptide: 8.8 – 4400 pg/mL; Proinsulin: 1.7 – 340 pg/mL; Des-31,32 proinsulin: 2.3 – 460 pg/mL; Des-64,65 proinsulin: 3.6 – 360 pg/mL.
